# Analysis of tracrRNAs reveals subgroup V2 of type V-K CAST systems

**DOI:** 10.1101/2025.03.07.641588

**Authors:** Marcus Ziemann, Alexander Mitrofanov, Richard Stöckl, Omer S. Alkhnbashi, Rolf Backofen, Wolfgang R. Hess

**Affiliations:** Faculty of Biology, Genetics and Experimental Bioinformatics, University of Freiburg, Freiburg, Germany; Chair of Bioinformatics, University of Freiburg, Freiburg, Germany; Institute of Microbiology & Archaea Centre, University of Regensburg, Regensburg, Germany; Center for Applied and Translational Genomics (CATG), Mohammed Bin Rashid University of Medicine and Health Sciences, Dubai Health, Dubai, United Arab Emirates; College of Medicine, Mohammed Bin Rashid University of Medicine and Health Sciences, Dubai Health, Dubai, United Arab Emirates; Signalling Research Centres BIOSS and CIBSS, University of Freiburg, Germany

**Keywords:** CRISPR, CRISPR-associated transposons, cyanobacteria, tracrRNA, transcriptional regulator

## Abstract

CRISPR-associated transposons (CAST) consist of an integration between certain class 1 or class 2 CRISPR-Cas systems and Tn7-like transposons. Class 2 type V-K CAST systems are restricted to cyanobacteria. Here, we identified a unique subgroup of type V-K systems through phylogenetic analysis, classified as V-K_V2. Subgroup V-K_V2 CAST systems are characterized by an alternative tracrRNA, the exclusive use of Arc_2-type transcriptional regulators, and distinct differences in TnsB and TnsC proteins. Although the occurrence of V-K_V2 CAST systems is restricted to Nostocales cyanobacteria, it shows signs of horizontal gene transfer, indicating its capability for genetic mobility. The predicted V-K_V2 tracrRNA secondary structure has been integrated into an updated version of the CRISPRtracrRNA program available on GitHub under https://github.com/BackofenLab/CRISPRtracrRNA/releases/tag/2.0.

## Introduction

Clustered regularly interspaced palindromic repeats and associated protein-coding genes (CRISPR-Cas systems) are adaptive immune systems in bacteria and archaea and the basis for the development of various genome editing tools (Hille *et al*. 2018; Makarova *et al*. 2020; Bharathkumar *et al*. 2022; Liu *et al*. 2022). Naturally occurring CRISPR-Cas systems act in three steps: adaptation, processing and interference (Makarova *et al*. 2015). The first step is initiated by contact with foreign DNA, like phage DNA, or plasmids. The cell can defend itself against this DNA and derive and store a short DNA fragment (spacer) from it in its own genome in an array consisting of spacers and short palindromic repeats (CRISPR array). In the second step, the CRISPR arrays are expressed as a long transcript, the pre-crRNA. This RNA will form specific hairpin structures based on the palindromic nature of the repeat sequences, which can be recognized by CRISPR-associated (Cas) proteins. The pre-crRNA then is processed into shorter CRISPR RNAs (crRNAs), which then form, together with other Cas proteins, the CRISPR-Cas interference complex. This complex is used, in the last step, to interact with DNA or RNA by base pairing. If the complex recognizes a sequence of sufficient similarity to the spacer, such as in the course of another phage attack, it cuts the target DNA or transcribed RNA and therefore protects the cell from infection.

CRISPR-Cas systems are widespread in most bacteria and archaea, but they do not share a common gene set or structure (Makarova *et al*. 2015, 2020). The systems are mainly classified into two classes, class 1, with multiple proteins forming the CRISPR- Cas effector complex, and class 2 using only a single effector complex protein. Both can be further divided into six types and over 30 different subtypes (Makarova *et al*. 2020). These subtypes differ in interference complex structure, composition and the nucleic acid type that is targeted (DNA or RNA). In this study, we focus on the subtype V-K, a class 2 CRISPR-Cas system integrated with a transposon (Strecker *et al*. 2019; Rybarski *et al*. 2021; Ziemann *et al*. 2023). This CRISPR-associated transposon (CAST) exists exclusively in cyanobacteria and is associated with the transposase genes *tnsB*, *tnsC* and *tniQ.* The proteins encoded by these genes facilitate genetic mobility by targeting tRNA genes using the CRISPR-Cas interference complex for the targeting mechanism (Koonin and Makarova 2024). The subtype V-K CRISPR-Cas components are minimal, with Cas12k as the only interference protein, a rather short CRISPR array (∼4 spacers per array) and a trans-activating crRNA (tracrRNA) forming the CRISPR-Cas complex. Recently, another related family of Tn7-like transposons was described that is targeting CRISPR arrays instead of tRNA genes. Also this family exists exclusively in certain cyanobacteria (Chacon Machado and Peters 2025).

The tracrRNA is a common element of class 2 CRISPR-Cas systems and is an adaptation unit between the crRNA and the Cas protein (Deltcheva *et al*. 2011). The crRNA can bind with its repeat region a specific anti-repeat region (AR) on the tracrRNA via base-pairing, while the Cas-protein binds the tracrRNA. In V-K systems, the location of these tracrRNA genes is also remarkably conserved downstream of *cas12k* and upstream of the typical type V-K CRISPR array, transcribed in the same direction (Strecker *et al*. 2019; Saito *et al*. 2021; Ziemann *et al*. 2023). The structure of type V-K tracrRNAs was established by two analyses of the *Scytonema hofmanni* CAST system (Xiao *et al*. 2021; Schmitz *et al*. 2022). These analyses revealed important RNA secondary structures such as the stem-loops P1-8, a pseudoknot, and repeat-antirepeat duplexes (Xiao *et al*. 2021) and provided a general three-dimensional structure for these types of tracrRNA. Most importantly, they showed the interaction with crRNA, which is here not facilitated by one, but two anti-repeat regions.

The first region is part of an RNA-triplet complex inside the second stem-loop (P2) and the second is at the 3’-end of the tracrRNA, approximately 120 nt apart from each other. This is rather unusual but occurs also in other type V tracrRNAs, like in V-B, V- F1 and V-G systems (Liao and Beisel 2021).

In previous studies, we compared known tracrRNA structures of this system (Xiao *et al*. 2021; Schmitz *et al*. 2022) to our database of CAST systems in order to develop an algorithm, called CRISPRtracrRNA, for the detection of these RNA genes (Mitrofanov *et al*. 2022). Here, we extend this analysis. We have developed CRISPRtracrRNA version 2 (V2), and were able to find an alternative tracrRNA, which appears as a structural derivative of the previously established version 1 tracrRNA (V1), and occurs only in a phylogenetically distinct group of CAST systems. This, as well as the exclusive association with a gene encoding a particular type of regulator, indicates that this constitutes an isolated subgroup of CAST systems (type V-K_V2).

## Results and Discussion

### Alternative tracrRNA of CAST type V-K

In our previously established database of CAST systems (Ziemann *et al*. 2023), tracrRNAs were predicted in approximately 77% of CAST systems. These tracrRNAs were predicted to be transcribed from loci located downstream of *cas12k* genes, in the same direction and upstream of the CRISPR array (if an array was present). These sequences were used as input to develop the CRISPRtracrRNA algorithm (Mitrofanov *et al*. 2022).

However, the algorithm occasionally detected partial tracrRNAs in CAST systems without previously established tracrRNAs, at the expected position downstream of *cas12k*. These predicted tracrRNAs could represent potentially truncated forms. However, the respective sequences between the *cas12k* genes and the CRISPR array are highly conserved (**Fig. 1B**). The previous genome-wide mapping of transcription start sites in *Anabaena* sp. PCC 7120 (Mitschke *et al*. 2011) indicated transcription of such a putative tracrRNA starting from position 3284143f on the forward strand (GenBank accession BA000019.2). This particular tracrRNA seemed to be degraded by the integration of a transposon. However, sequence comparison showed that the associated promoter region is conserved (**Fig. 1B**). The respective transcription start site was assigned as the start of the alternative tracrRNA, here called tracrRNA_V2.

**Figure 1.**
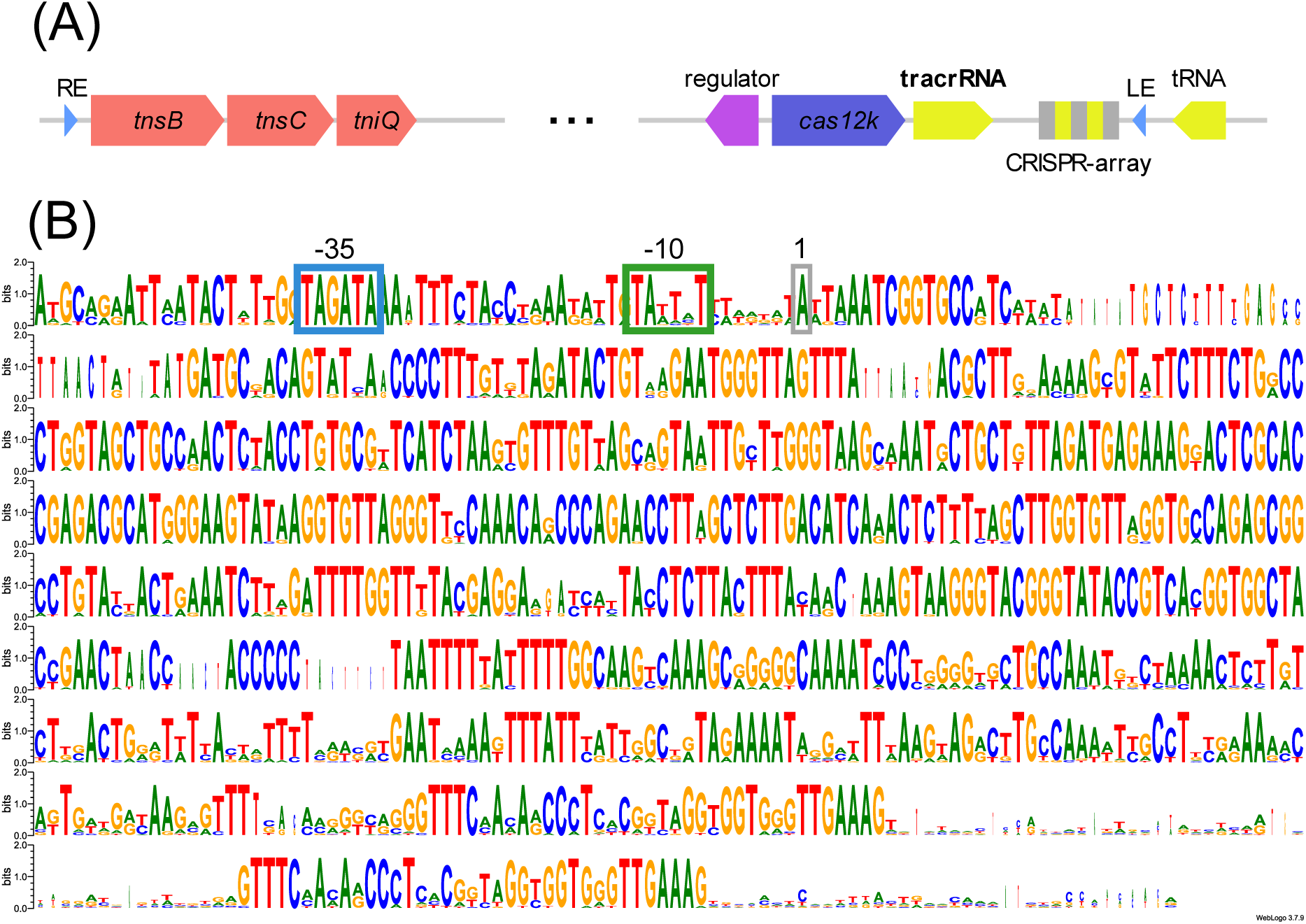
Gene arrangement in type V-K CAST systems and the conservation of V2 tracrRNAs. **(A)** The gene arrangement, shows the CAST system from the right (RE) to the left end (LE). The repeat-spacer array is drawn in gray and yellow, other colors indicate gene functions (red, transposase genes; rose, CAST regulators; dark purple, *cas12k* effectors; yellow, tRNAs and tracrRNAs). The scheme is not drawn to scale. **(B)** Sequence alignment of the areas between *cas12k* and the CRISPR array, based on 19 systems with V2 tracrRNAs and the previously established promoter sequence of *Anabaena* sp. PCC7120 (Mitschke *et al*. 2011). Conserved promoter elements are boxed in blue (−35 element), green (−10 element), and gray (transcription start site). The individual sequences can be found in supplemental **Dataset 1**. The transcription start site (position +1) was experimentally mapped in *Anabaena* (*Nostoc*) sp. PCC 7120 (Mitschke *et al*. 2011) and taken as reference for the other sequences.

An additional search for the *cas12k*-gene and tracrRNA-like sequences resulted in the identification of ten additional CAST systems with tracrRNA_V2, which increased our dataset to 128 type V-K CAST systems in total (**Dataset 2** and **3**). Further analysis showed that these V2 systems were all closely related. They are exclusively associated with regulatory genes encoding the Arc_2 regulators (Ziemann *et al*. 2023), and the phylogenetic analysis showed their relatedness in a single, coherent clade (**Dataset 2; Fig. 2**). Phylogenetic analyses of the associated transposases yielded a similar tight clade (**Fig. S1-S3**), further suggesting that these CAST systems are genetically closely related and distinct from other systems.

**Figure 2.**
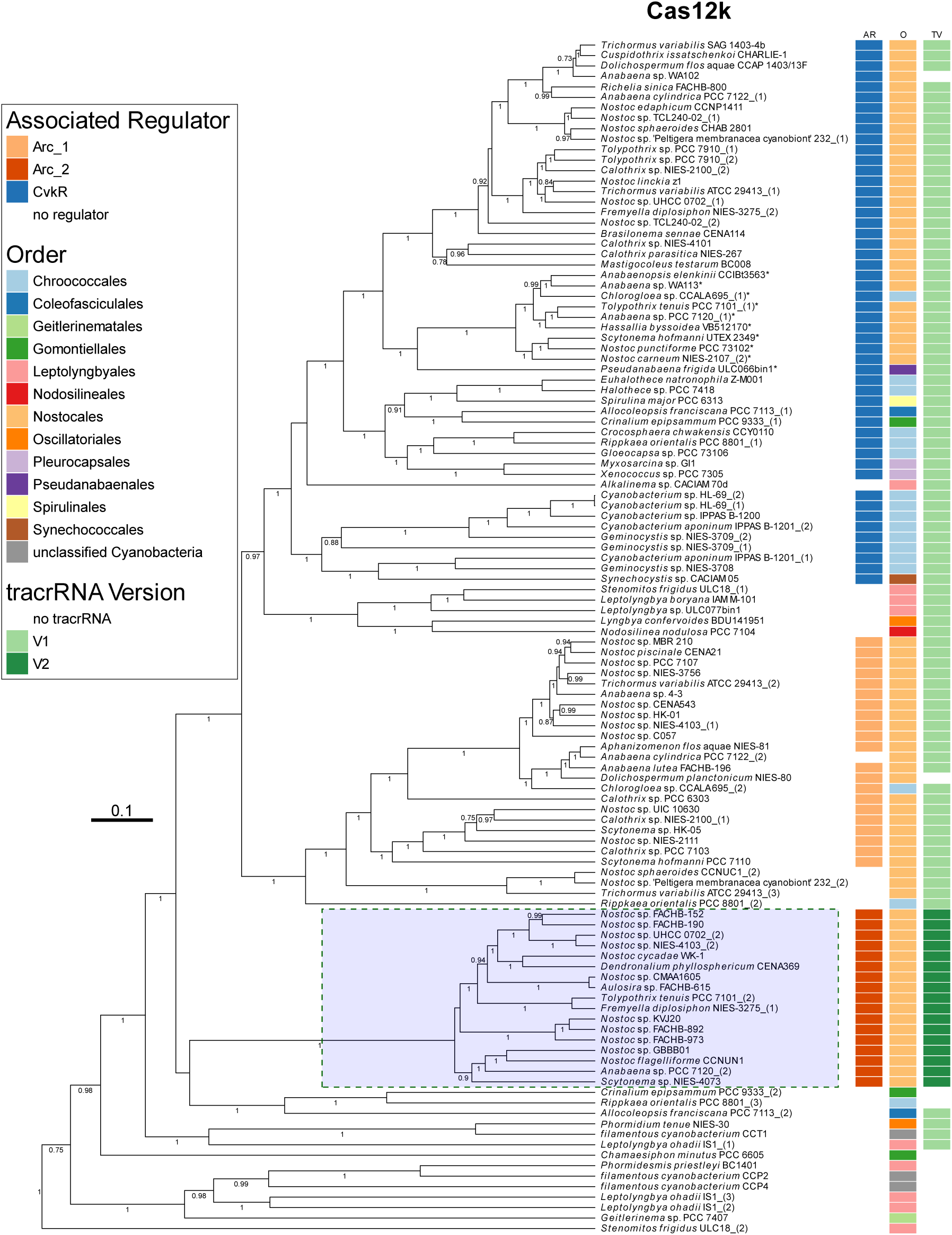
Phylogenetic tree of CAST systems based on Cas12k sequences, labeled by the respective strain names. The distinct group of type VK-2 CAST systems is shaded in blue. A selected group of V1 CAST systems are marked with asterisks. These systems were later used for comparisons with V2 systems. Numbers at branches indicate respective posterior probabilities. The respective associated regulators (AR), taxonomic order (O) and tracr types (TV) are indicated as given. The protein sequences were aligned with M-coffee (Di Tommaso *et al*. 2011) and analyzed by BEAST (Suchard *et al*. 2018). For the multiple sequence alignment of Cas12k proteins of all V2 and 10 selected V1 CAST systems (here marked with asterisks), see **Fig. S7**.

Based on these evolutionary connections, the 21 V2-associated systems were further analyzed. The DNA fragments between the *cas12k* gene and the end of the CRISPR array were aligned and used for an analysis by shapes studios (Janssen and Giegerich 2015), a webtool application to predict possible RNA structures out of sequence alignments. For this, only 19 of these sequences were used because the other two showed signs of degradation (see **Dataset 1** excluding the promoter sequence from *Anabaena* sp. PCC 7120; **Fig. 1B**). The potential structures or shapes from this analysis were then investigated to find common base pairings that existed in over 50% of the structures. This approach, together with the established structures of the original tracrRNA V1, was used to predict a potential V2 tracrRNA structure (**Fig. 3, S5** and **Dataset 1** and **5**). For comparison, the same method was used with ten other sequences, which based on the phylogeny of Cas12k, were V1 tracrRNAs including those from the CAST systems of *Anabaena* sp. PCC 7120 and *S. hofmanni* (**Fig. 2, S5** and **Dataset 4** and **6**).

**Figure 3.**
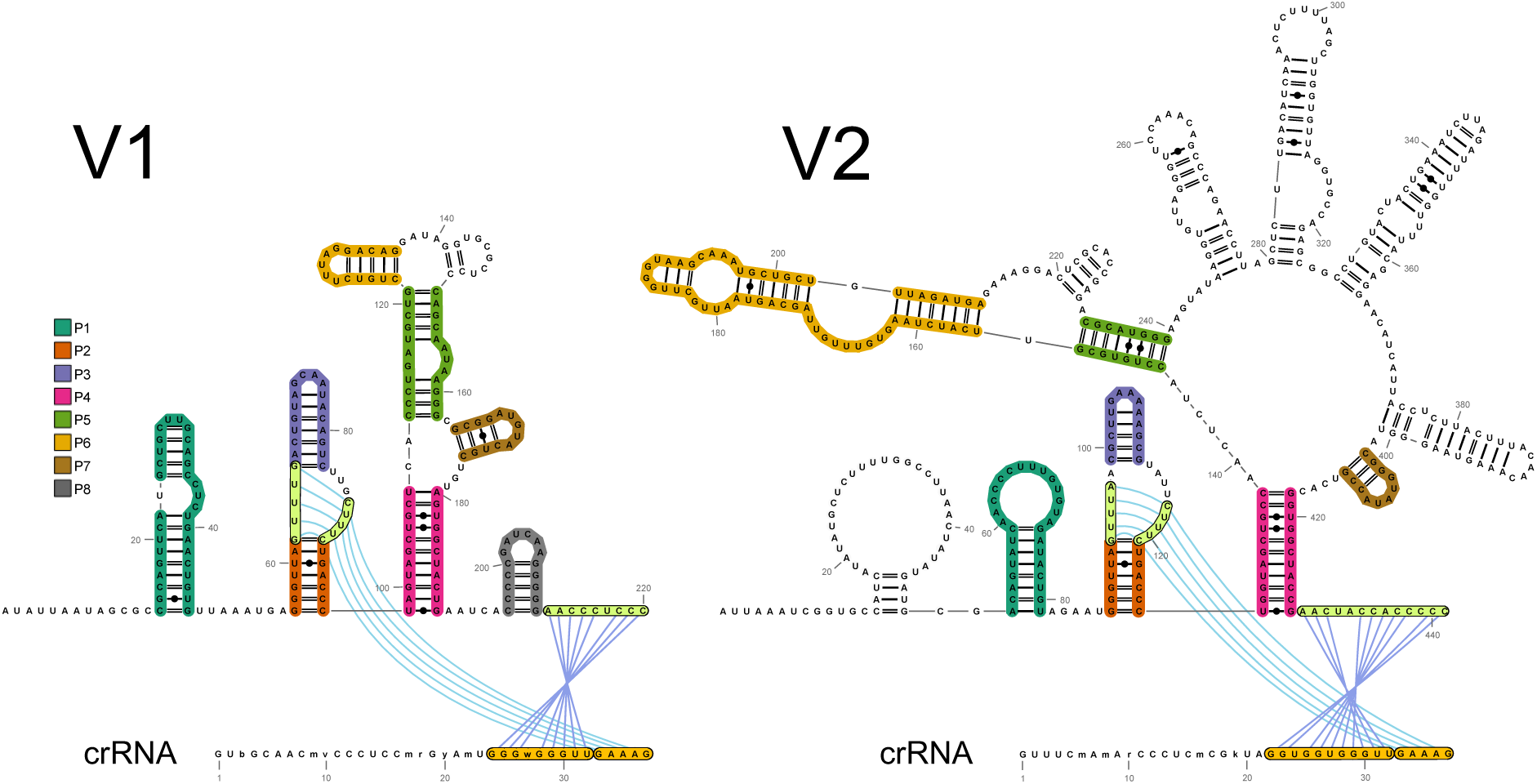
Predicted shapes of tracrRNAs V1 and V2. The individual stem-loops are marked with different colors and anti-repeat regions are marked in yellow. The structures were established by shapes studio (Janssen and Giegerich 2015) and based on the alignments of 10 selected V1 systems and 19 V2 systems (see Fig. 2 and **Datasets 4–6**). Gaps were deleted from the structure. The consensus sequence of the respective crRNA repeat was added and the tracrRNA-bound segments are marked in orange. Blue lines indicate potential interactions between individual nucleotides. The figure was drawn with RNAcanvas (Johnson and Simon 2023). See **Fig. S4** for sequence alignments and **Fig. S5** for further context.

The predictions indicate that 7 of the 8 stem-loops typical for type V-K V1 tracrRNAs (Xiao *et al*. 2021) exist in V-K V2 tracrRNAs as well (**Fig. 3**). Clear sequence and structural similarities exist for stem-loops P2-P7, while the stem-loop P1 structure differs. Remarkably, we identified four additional stem-loops that appear to be inserted between P5 and P7 (X1-4; **Fig. 3** and **S5**) in the V2 tracrRNA relative to the V1 tracrRNA.

Most importantly, the V2 tracrRNA includes the central triplex necessary for the tracrRNA-crRNA interaction (Anti-repeat 1) (Schmitz *et al*. 2022). Therefore, it can be assumed to bind the crRNA in the same manner as in V1 systems. However, the typical second Anti-repeat area (AACCCnCCC) is missing in V2. Instead, the structure prediction of Shapes Studio analysis and sequence comparisons indicated a different Anti-repeat 2 right after the P4 stem-loop (AACUACCACCCCC) (see **Fig. 3, S4, S5**; **Dataset 5**). This difference in the anti-repeat can also be seen in the CRISPR array repeats. The 3’-end, which in V1 binds the tracrRNA, changed from 5’- wkrGGyGGGTTGAAAG-3’ to 5’-**<u>GGT</u>**GG**<u>T</u>**GGGTTGAAAG-3’ in V2 (see **Fig. S4**). The tracrRNA-crRNA-binding in V2 seems to be established by three mismatching variable nucleotides, similar to the interaction in tracrRNA V1.

### In vivo proof of tracrRNA_V2 of CAST type V-K

To prove the existence of the tracrRNA_V2, we cultivated *Scytonema* sp. NIES-4073, which contains a complete CAST system with an Arc_2 repressor and a potential V2 tracrRNA. The previously used model organism *Anabaena* sp. PCC 7120 also contains a type V-K_V2 system. However, the tracrRNA was degraded by the insertion of a transposon inside its gene and, therefore, was insufficient for this analysis. The *Scytonema sp*. NIES-4073 strain was chosen because this CAST system showed no sign of degradation and because of its availability from the NIES strain collection. After 3 months of cultivation, RNA was extracted and used for northern blots against the tracrRNA and the CRISPR array. Several bands were detected for both transcripts, indicating their expression and maturation (**Fig. 4**). Two prominent RNA fragments were detected in the northern blot against the tracrRNA probe at 450 nt and 700 nt. The 450 nt fragment is likely the mature tracrRNA. Interestingly, a 700 nt band was also observed in the northern blot of the CRISPR array, which indicates that tracrRNA and CRISPR array are transcribed into a joint precursor.

**Figure 4.**
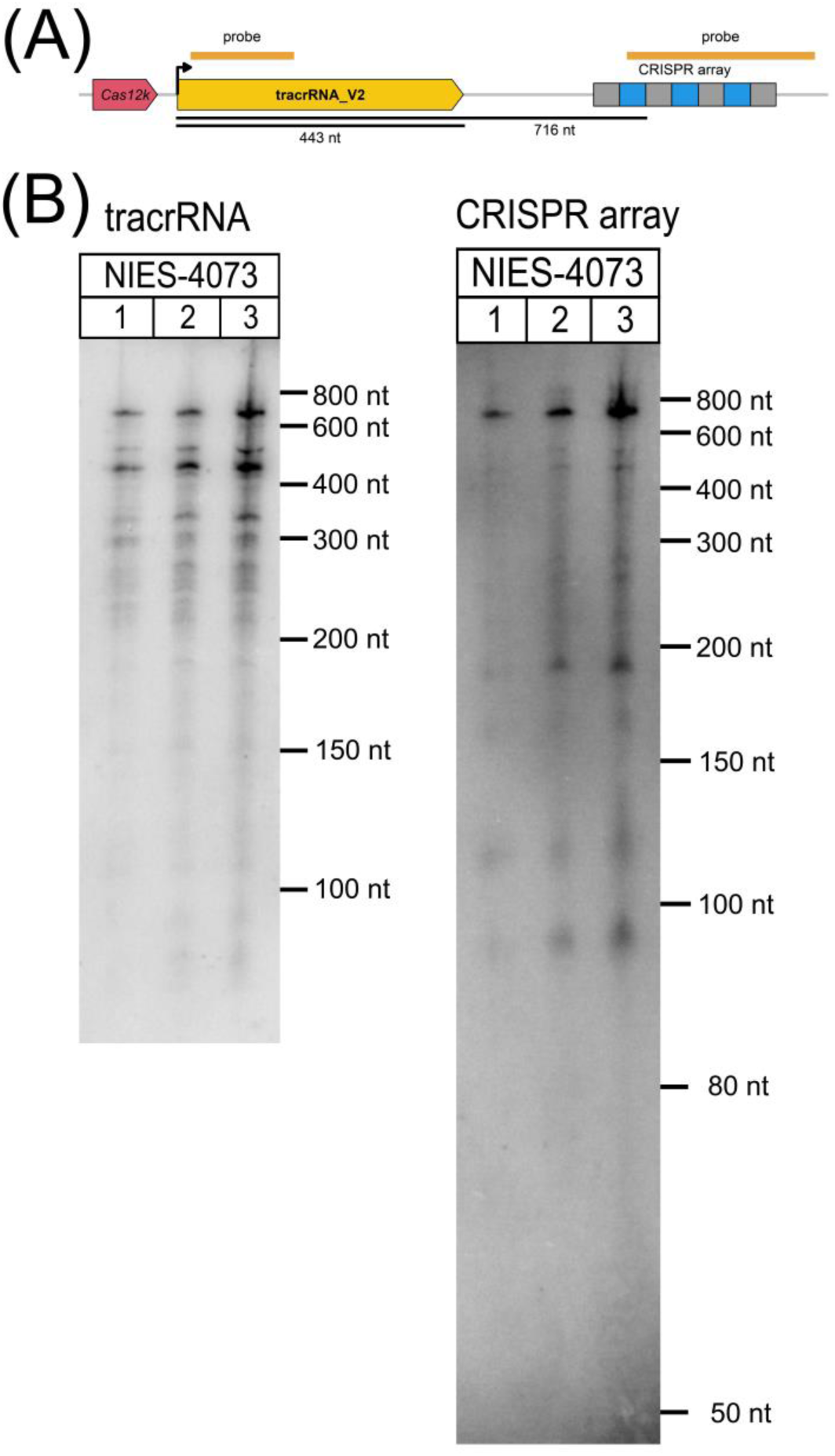
Experimental detection of the V2 tracrRNA. **(A)** Location of the tracrRNA and CRISPR array in *Scytonema sp.* NIES-4073 with marked positions of northern blot probes and transcription initiation site (bend arrow). The figure is not drawn to scale. **(B)** Northern blot hybridization against probes for tracrRNA and the CRISPR array.

### Phylogenetic analysis of CAST systems

To better understand the characteristics of the type V-K V2 CAST systems described here, phylogenetic analyses were performed for the most prominent CAST proteins, TnsB, TnsC, TniQ (**Fig. S1, S2 and S3**) and Cas12k (**Fig. 2**). All four phylogenetic trees placed systems with a V2 tracrRNA into a distinct group among all CAST systems. The analysis of the V2 Cas12k proteins showed with a minimal shared sequence identity of 60.2% their closer relatedness to the exclusion of all other Cas12k proteins (shared identity ≤47.5% between V1 and V2) (**Fig. 2, S7**), but very few structural differences compared to other Cas12k proteins. This is in contrast to TnsB (shared sequence identity in V2 ≥66.0% and ≤43.1% between V1 and V2), where V2 systems showed a difference in domain organization compared to the homologs from V1 systems. Based on the MSA and predictions by Alphafold 3 (Abramson *et al*. 2024), the specific DNA-binding region lγ is 20 amino acids larger, forming a plateau of three additional helices (Park *et al*. 2022)(**Fig. S6**, **Dataset 7** and **8**). This change enlarges the domain significantly, which is known to bind the IS elements of the CAST transposon during the transposition (Park *et al*. 2022).

However, the Alphafold prediction showed no interaction of the extra region with DNA (**Fig. S6**, **Dataset 8**). Moreover, the alignment of the IS elements indicated similar lengths for the long terminal repeats of V1 and V2 systems (**Fig. 5B**). Interestingly, the alignments also showed unique consensus sequences for V2 CAST systems with higher levels of conservation, consistent with the higher conservation of TnsB. In addition, the V2 systems also showed a very consistent insertion distance to its anchor protospacer (Ziemann *et al*. 2023). The distance between PAM sequence and LE element is 54 nt, shorter than the average distance of 62 nt in V1 type V-K systems, which was also predicted for the system in *S. hofmanni* (Strecker *et al*. 2019)(**Fig. 5A**).

**Figure 5.**
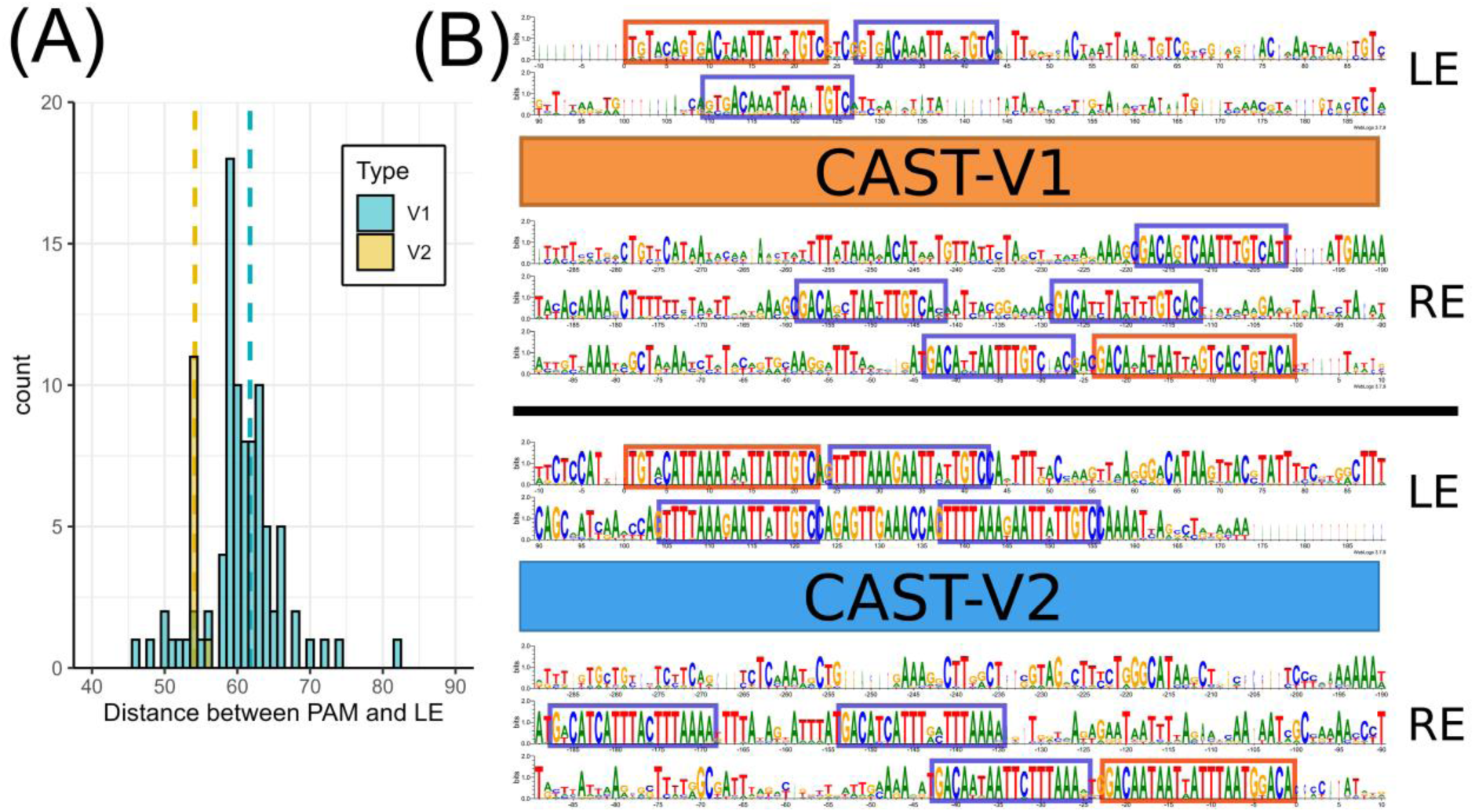
Comparison between VK_V1 and V2 systems in insertion distance and IS elements. **(A)** Distance between the PAM sequence of the anchor-protospacer and the LE element in V1 and V2 systems in comparison. The dashed lines show the average distances. Note, that there is one outlier (V1) with a distance 146 nt that was not included. **(B)** Alignment of identified IS elements of selected V1 and all V2 CAST systems. Ten related V1 systems were used (see strains marked by asterisks in Fig. 2 and in **Dataset 2**). The long terminal repeats (LTR) and short repeats (SR) of the sequences were marked in orange (LTR) and blue (SR) boxes.

The other V2 transposases showed fewer differences than their V1 homologs. The V2 TnsC showed distinct prolongations of the C-terminal and N-terminal domains compared to V1 TnsC (shared sequence identity in V2 ≥79.1% and ≤33.8% between V1 and V2). The N-terminal domain is important for the interaction with TniQ and the C-terminal domain interacts with TnsB in the transposition complex (Tenjo-Castaño *et al*. 2024). For this reason, V2 TnsC proteins are also significantly larger (306 aa) than V1 TnsC (278 aa); however, this does not seem to be related to an additional domain. TniQ V2 is highly conserved and lacks differences in domain arrangements between V1 and V2 V-K systems (shared sequence identity in V2 ≥84.1% and ≤61.1% between V1 and V2).

### An updated version of the CRISPRtracrRNA algorithm

The CRISPRtracrRNA tool was originally designed to facilitate the efficient identification of tracrRNA sequences in genomes containing Type II and Type V CRISPR-Cas systems (Mitrofanov *et al*. 2022). The algorithm integrates multiple methodologies to robustly detect tracrRNA sequences. These methodologies include CRISPRidentify (Mitrofanov *et al*. 2021) and CRISPRcasIdentifier (Padilha *et al*. 2020) for identifying CRISPR arrays and classifying associated *cas* genes, terminator sequences, and perform sequence-structure analysis using a pre-trained covariance model. The accuracy of the covariance model is highly dependent on both the quality and size of the training dataset. Therefore, the here described V2 tracrRNA was added to our database.

Previously, the type V covariance model was trained on 91 sequences. To enhance its quality, we incorporated 19 additional novel sequences into the dataset. Of these, 10 sequences were added to the training set, while the remaining 9 were retained as a test set to validate model performance. Model training was conducted using the GraphClust2 Galaxy pipeline (Afgan *et al*. 2018; Miladi *et al*. 2019; Galaxy Community 2024), consistent with our prior methodology.

Upon training the updated model, we observed the emergence of four distinct clusters, compared to the two clusters in the previous version. This prompted a thorough performance evaluation of the new model. The calibration of the covariance model was carried out using the cmcalibrate module of the Infernal tool (Nawrocki and Eddy 2013). This calibration process is critical for optimizing model parameters, including E-value thresholds, which are essential for assessing statistical significance in sequence search results.

Following calibration, we evaluated the model’s performance on both the training and test datasets using cmscan module of the Infernal tool (Nawrocki and Eddy 2013). The results demonstrated a significant improvement in performance on the test set, without any compromise in training set performance (**Fig. S8**). Additionally, a comparative analysis of sequence coverage between the updated and previous models revealed that the increased number of clusters in the new model provided substantially improved coverage across both the training and test datasets (**Fig. S9**).

The updated model has been integrated into the CRISPRtracrRNA pipeline, now released as CRISPRtracrRNA 2.0. In this release, we also resolved compatibility issues with the Conda environment that had accumulated over time. The updated tool is now available on our GitHub repository, providing a robust and enhanced framework for tracrRNA sequence identification.

## Conclusion

Cyanobacteria are a rich source of CRISPR-Cas systems (Scholz *et al*. 2013; Behler *et al*. 2018; Hou *et al*. 2019; McBride *et al*. 2020; Reimann *et al*. 2020). Among these systems are the type V-K CAST systems that are unique to Nostocales cyanobacteria (Strecker *et al*. 2019; Rybarski *et al*. 2021; Ziemann *et al*. 2023). Here, we show that the here described type V-K_V2 systems are set apart by their tracrRNAs, the family of the associated transcriptional regulator, and distinct features of the associated Cas12K, TnsB and TnsC proteins. The newly identified V2 tracrRNA sequences were incorporated into the training dataset for the CRISPRtracrRNA algorithm, which has been extended and updated accordingly.

## Materials and methods

### Cultivation of cyanobacteria and RNA extraction

The strain *Scytonema* sp. NIES-4073 was ordered by the NIES-collection (Microbial Culture Collection at the National Institute for Environmental Studies) and grown in liquid MDM-medium (Watanabe 1960), under permanent white light illumination of 10-15 μmon photons m^−2^ s^−1^ at 20 °C. The cultivation was done exclusively in cell culture flasks without shaking. Because of the slow growth new medium was added regularly. *Scytonema* sp. NIES-4073 forms thick cell clusters in liquid medium, which made a measurement of optical density impossible. 500 mL Flasks were filled with 150 mL MDM-medium and inoculated with NIES-4073 strain. Fresh medium was added in regular intervals. After 3 months, 500 mL cultures were harvested by filtering (0.85 µm). The cells were transferred to screw cap tubes via resuspension in 300 μL MDM- medium (Watanabe 1960), together with 500 μL glass beads (250 μL 0.1–0.25 μm in diameter and 250 μL 0.25-0.5 μm in diameter). 500 μL PGTX(Pinto *et al*. 2009) was added and the tubes were frozen in liquid nitrogen. Cell disruption (3 cycles 30 Hz for 10 minutes with short breaks in between) with mixer mill MM400 (Retsch) was performed at 4 °C. Samples were separated from the beads and incubated for 30 min at 65 °C in a water bath, one volume chloroform:isoamyl alcohol (24:1) was added, and samples were incubated for 10 min at room temperature with several vortexing cycles in between. After centrifugation for 3 min at 3250 g in a swing-out rotor, the supernatant was transferred to a fresh tube, and one volume of chloroform:isoamyl alcohol was added. This step was repeated twice. RNA was precipitated by the addition of one volume of isopropanol.

Eight μg of total RNA per sample were analyzed by northern hybridization using single-stranded radioactively labeled RNA probes as described (Ziemann *et al*. 2023). The probes were generated *in vitro* from PCR-generated templates (see Supplementary **Table S1** for primers).

### Phylogenetic analysis of Cas12k and the transposases

The identified Cas12k, TnsB, TnsC and TniQ proteins were compared to each other and analyzed for alternative start positions to correct potential incorrect annotations, as described in (Ziemann *et al*. 2023). The used sequences deviating from the original annotations are noted in the **Dataset 2**. Proteins with truncated sequences were removed from the analysis. The sequences were aligned by M-coffee (Notredame, Higgins and Heringa 2000; Di Tommaso *et al*. 2011) and further analyzed by the BEAST algorithm (v2.7.7) (Suchard *et al*. 2018: 1; Bouckaert *et al*. 2019). The phylogenetic analyses were calculated with a strict clock model and Yule model as tree prior, using the standard parameters (birthRate.t: Uniform[0,0, Infinity]; initial = [1.0][0.0, ∞]). The MCMC chain was sampled for TniQ and Cas12k every 1000 steps over 5 million generations (Yule 1925; Henikoff and Henikoff 1992; Gernhard 2008) and every 1000 steps over 1 million generations for the analysis of TnsC and TnsB. In order to ensure a reasonable effective sample size (ESS; threshold at 200), the results were analyzed by the Tracer v 1.7.2 tool (Rambaut *et al*. 2018) and the first 10% of trees were discarded from the analysis. The trees were then generated by the tree annotator tool (from the BEAST package).

### tracrRNA predictions

Homologous sequences (see **Dataset 1** for the precise sequences) between *cas12k* and the second repeat of the CRISPR array were aligned by M-coffee (Notredame, Higgins and Heringa 2000; Di Tommaso *et al*. 2011) and used as input for the webtool RNAalishapes from the Shapes Studio server (Janssen and Giegerich 2015). The resulting structures were then analyzed to identify nucleotide binding pairs that exist in more than 50% of all predicted structures. This analysis was done with 19 V2 sequences and 10 selected V1 systems for comparison (see **Fig. 3**, **Dataset 1, 4**). The resulting structures were then compared to the known V1 tracrRNA structures and stem-loops. For the model shown in **Fig. 3**, gaps were removed from the structure to simplify the visualization. The complete structures are available in supplement **Fig. S5** and **Dataset 5 and 6**. The specific detected stem-loops were also aligned and compared to each other (**Fig. S4**) using the msa-package from R and the webtool weblogo (Schneider and Stephens 1990; Crooks *et al*. 2004; Bodenhofer *et al*. 2015).

