## Supplemental Information for "Analysis of tracrRNAs reveals subgroup V2 of type V-K CAST systems"

### Supplementary Information

**Overview on Supplementary Datasets:** p. 2

**Supplementary Figures:** p. 3

**Supplementary Table S1:** p. 12

**Supplementary References:** p. 12

### Overview on Supplementary Datasets

**Dataset 1:** Sequences of 19 V2 CAST systems from 100 nt upstream of the tracrRNA TSS to 100 nt downstream of the first repeat of the CRISPR array and in addition the promoter sequence of *Anabaena* sp. PCC7120 [3284143f] 100 nt upstream to TSS and 60 nt downstream of TSS. All identified V2 systems were used in our analyses, except two systems with degraded tracrRNAs. All sequences are named after the corresponding Cas12k protein of the CAST system. Provided as separate file in FastA format.

**Dataset 2:** Available data on all known CAST systems. The table includes, among others, sequences and positions of *cas12k*, *tnsB*, *tnsC*, *tniQ* and regulator proteins. The “start”, “end” and “border” values in the dataset correspond to the position on the related sequence in Dataset 2. Provided as separate Excel file.

**Dataset 3:** All sequences of known CAST systems, including the surrounding regions. The sequences were named after the corresponding Cas12k protein of the CAST system. Provided as separate file in FastA format.

**Dataset 4:** Sequence of 10 V1 CAST systems from 100 nt upstream of tracrRNA to 100 nt downstream of the first repeat of the CRISPR array. The systems were chosen from closely related V1 systems including already established CAST systems from *Anabaena sp*. PCC 7120 and *Scytonema hofma*nni UTEX 2349. All sequences are named after the corresponding Cas12k protein of the CAST system. Provided as separate file in FastA format.

**Dataset 5:** Structure predictions of tracrRNA V2 from Shape studios RNAalishapes program (Janssen and Giegerich 2015) based on the alignment of the sequences from Dataset 1, excluding the promoter sequence of of *Anabaena sp*. PCC7120. Provided as separate txt file.

**Dataset 6:** Structure predictions of tracrRNA V1 from Shape studios RNAalishapes program (Janssen and Giegerich 2015) based on the alignment of the sequences from Dataset 4. Provided as separate txt file.

**Dataset 7:** Alphafold 3 (Abramson *et al.* 2024) prediction of type V1 TnsB protein [WP_084763316.1] and DNA, based on the sequence surrounding the *Scytonema hofma*nni UTEX 2349 CAST system. Provided as separate file in CXS format.

**Dataset 8:** Alphafold 3 (Abramson *et al.* 2024) prediction of type V2 TnsB protein [BAZ25524.1] and DNA, based on the sequence surrounding the *Scytonema sp.* NIES-4073 CAST system. Provided as separate file in CXS format.

### Supplementary Figures

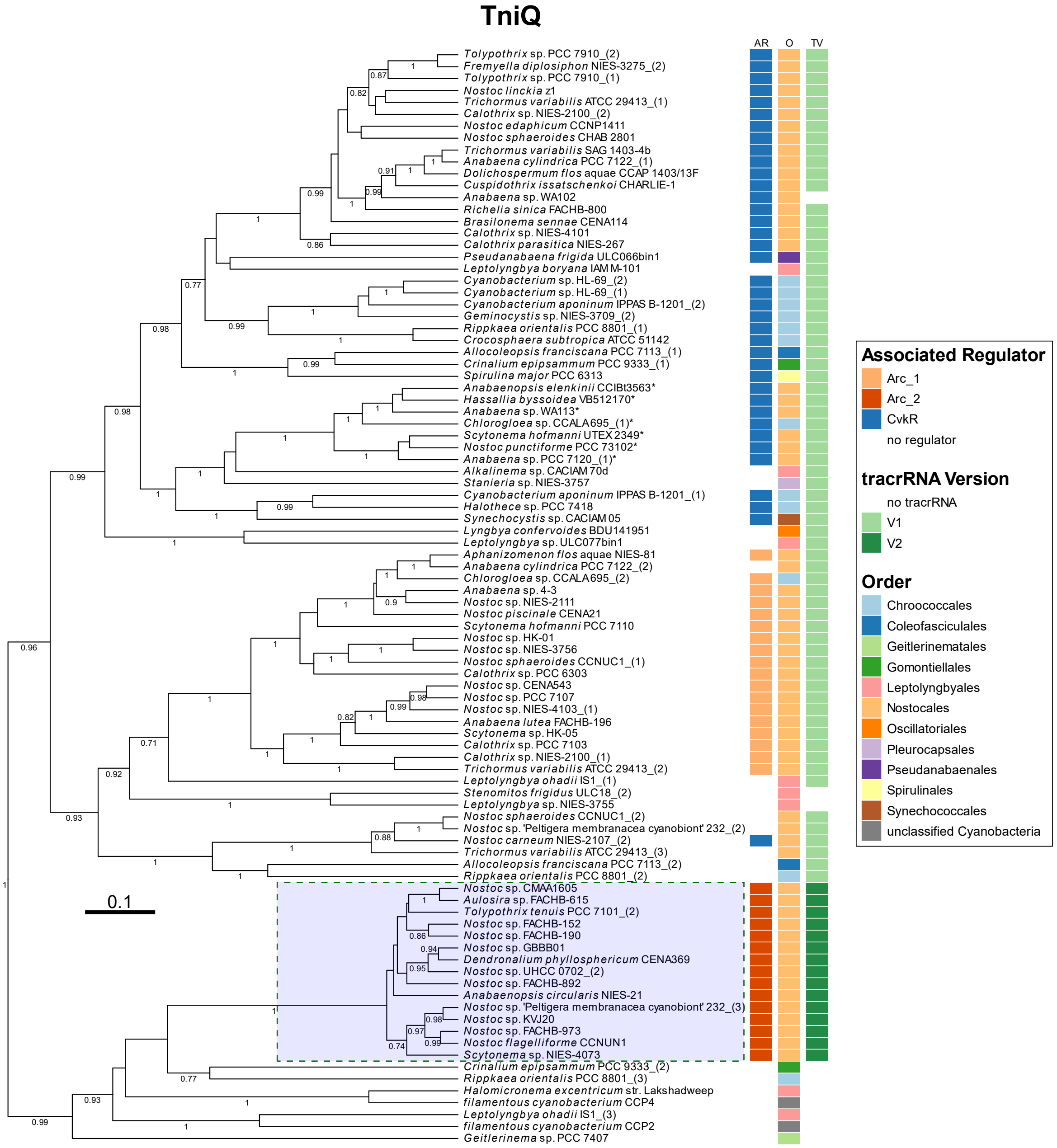


**Supplemental Figure S1. Phylogenetic tree of CAST systems based on TniQ sequences, labeled by the respective strain names.** The distinct group of type VK-2 CAST systems is shaded in blue. A selected group of V1 TniQ proteins are marked with asterisks. These proteins were later used for comparisons with V2 systems. Numbers at branches indicate respective posterior probabilities. The respective associated regulators (AR), taxonomic order (O) and tracr types (TV) are indicated as given. The protein sequences were aligned with M-coffee (Di Tommaso et al. 2011) and analyzed by BEAST (Suchard et al. 2018). For the multiple sequence alignment of TniQ proteins (all V2 and 10 selected V1 CAST systems, here marked with an asterisk), see **Fig. S7**.

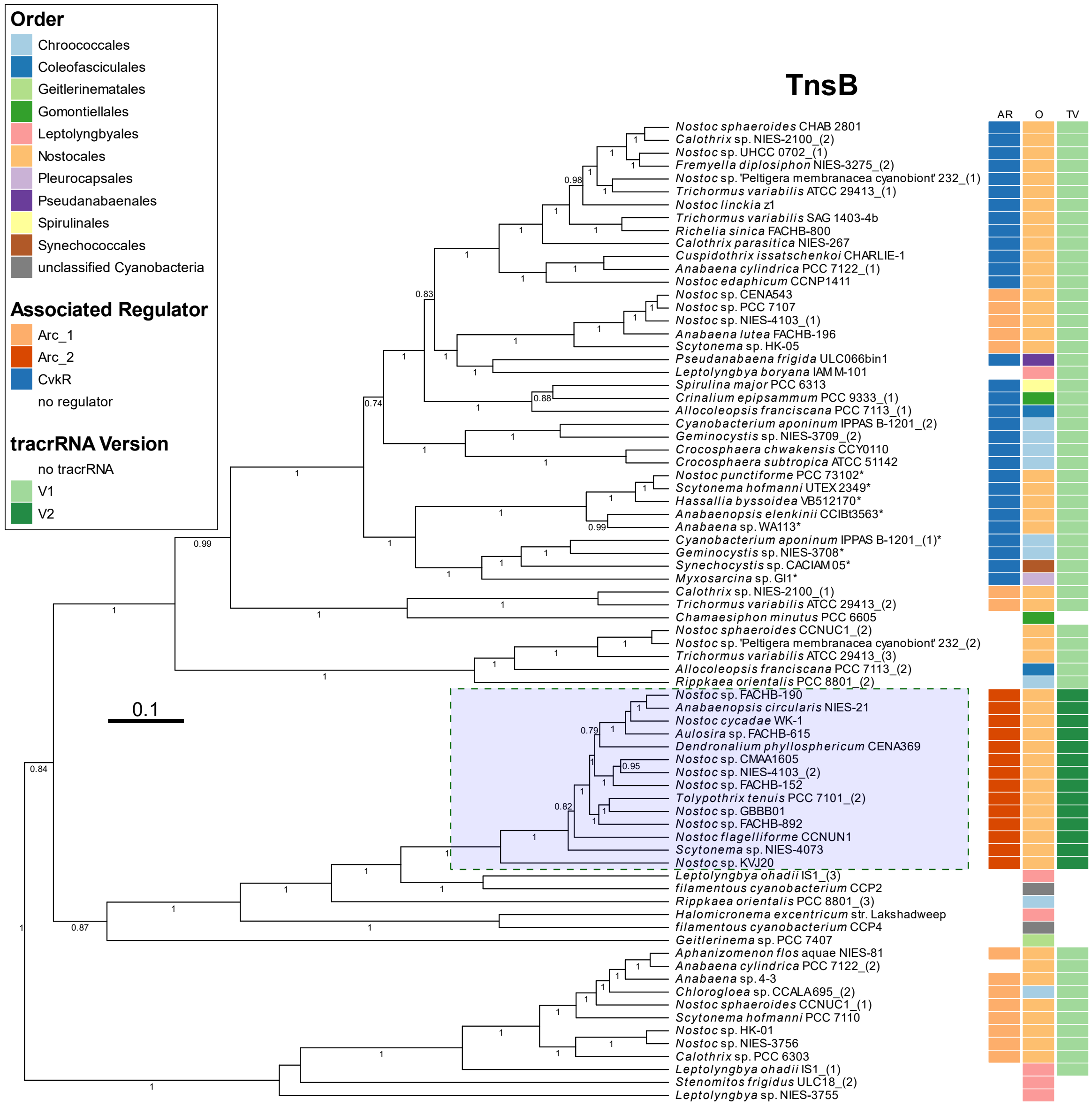


**Supplemental Figure S2. Phylogenetic tree of CAST systems based on TnsB sequences, labeled by the respective strain names.** The distinct group of type VK-2 CAST systems is shaded in blue. A selected group of V1 TnsB proteins are marked with asterisks. These proteins were later used for comparisons with V2 systems. Numbers at branches indicate respective posterior probabilities. The respective associated regulators (AR), taxonomic order (O) and tracr types (TV) are indicated as given. The protein sequences were aligned with M-coffee (Di Tommaso et al. 2011) and analyzed by BEAST (Suchard et al. 2018). See **Fig. S6** for the TnsB multiple sequence alignment and an Alphafold prediction of the N-terminal structures.

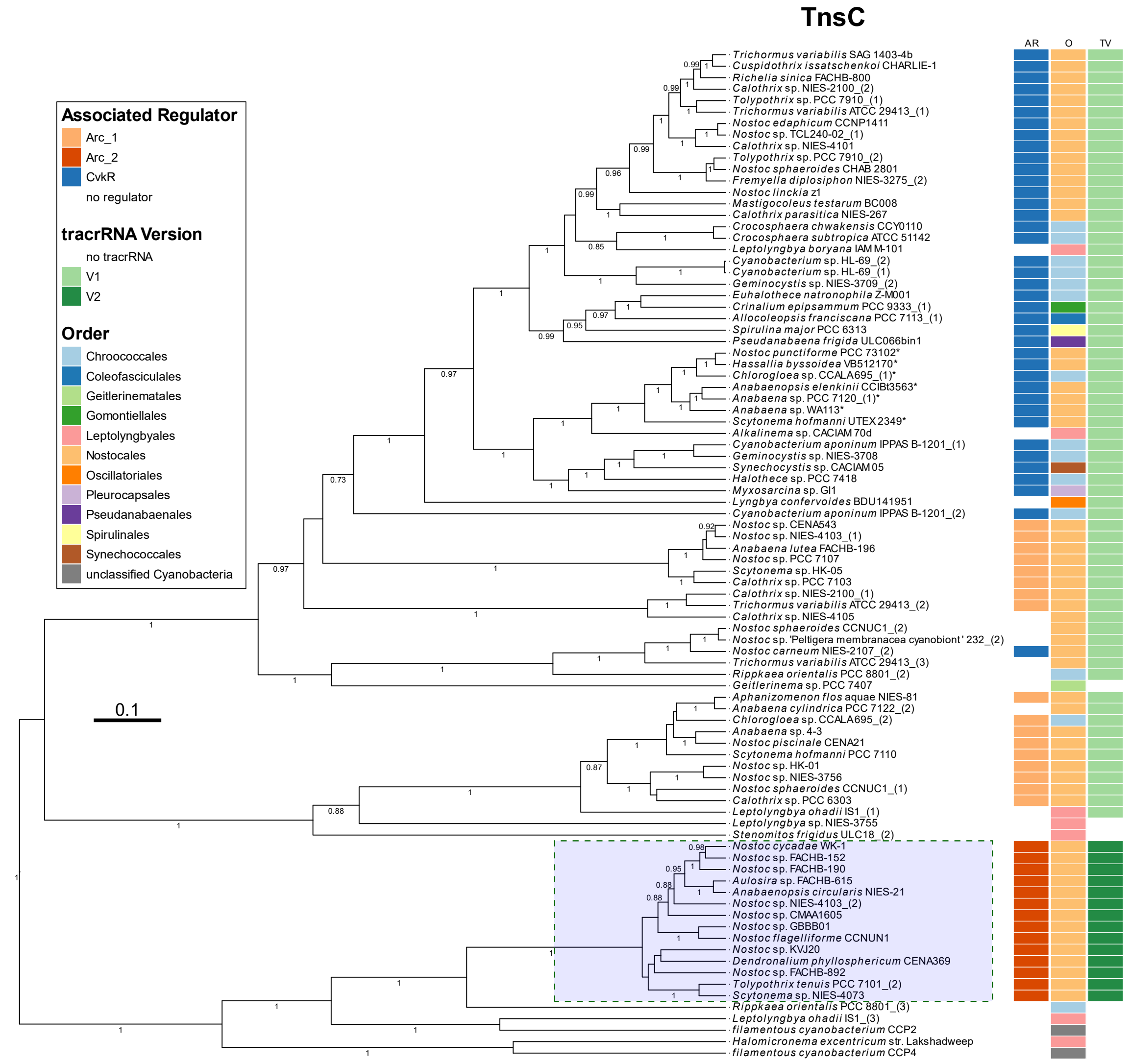


**Supplemental Figure S3. Phylogenetic tree of CAST systems based on TnsC sequences, labeled by the respective strain names.** The distinct group of type VK-2 CAST systems is shaded in blue. A selected group of V1 TnsC proteins are marked with asterisks. These proteins were later used for comparisons with V2 systems. Numbers at branches indicate respective posterior probabilities. The respective associated regulators (AR), taxonomic order (O) and tracr types (TV) are indicated as given. The protein sequences were aligned with M-coffee (Di Tommaso et al. 2011) and analyzed by BEAST (Suchard et al. 2018). For the multiple sequence alignment of TnsC proteins (all V2 and 10 selected V1 CAST systems, here marked with an asterisk), see **Fig. S7**.

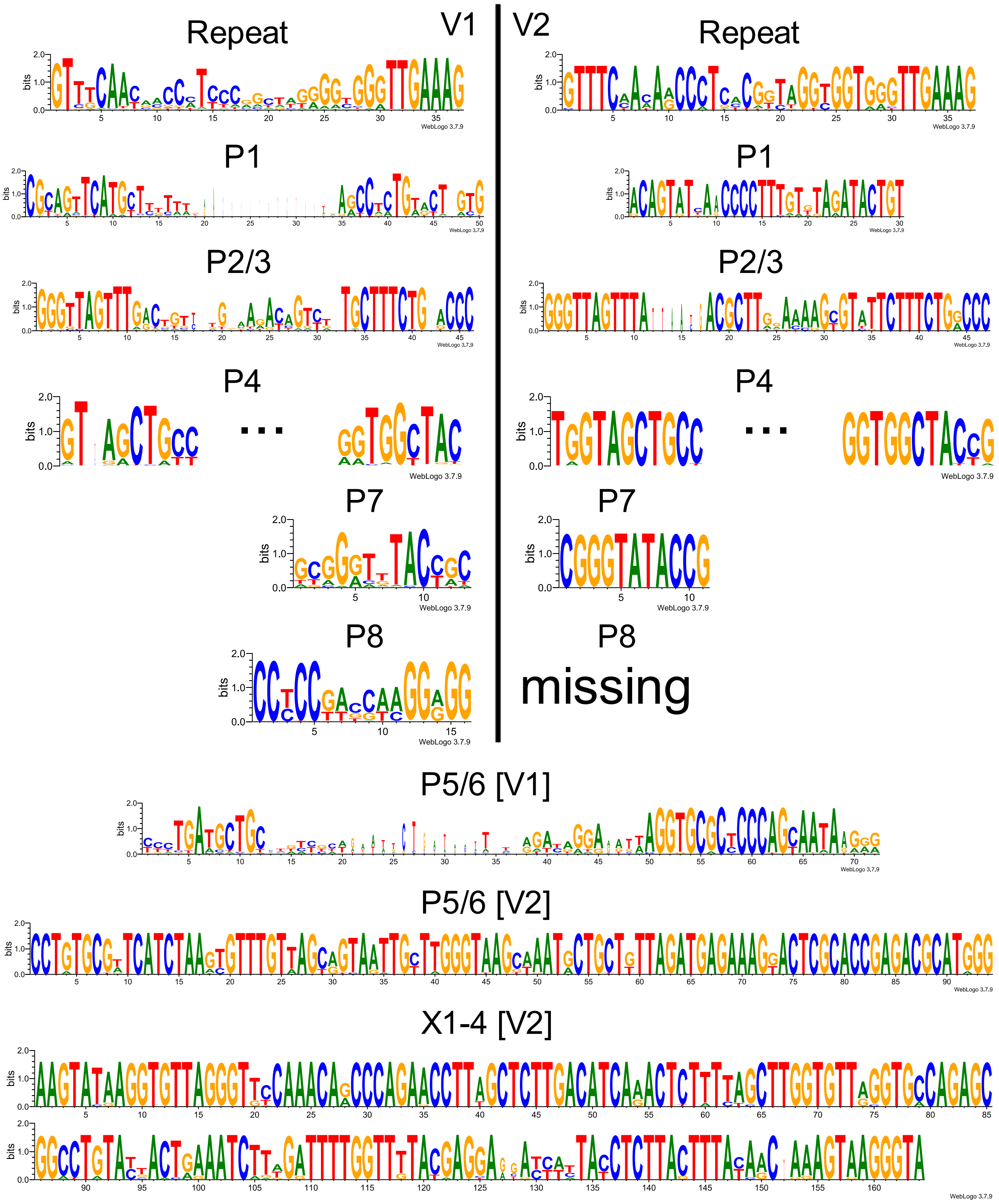


**Supplemental Figure S4**. **Sequence alignment and comparison between V1 and V2 tracrRNA and repeat sequences.** The sequences of all available V1 and V2 systems were compared to each other based on their specific stem-loops (P1-8) and CRISPR array repeats. This figure extends **Fig. 3**.

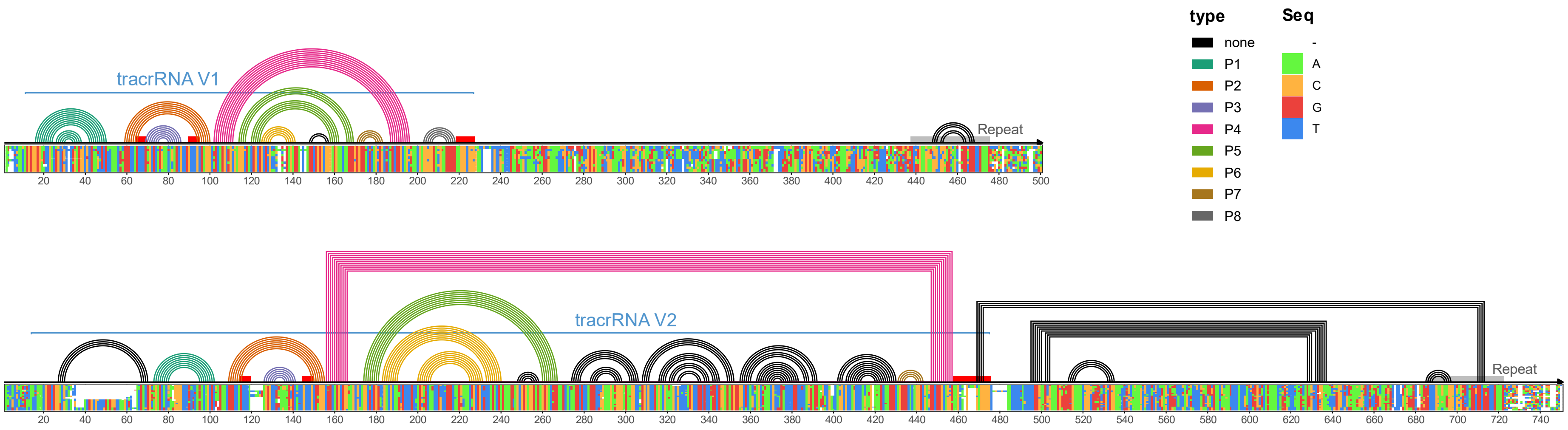


**Supplemental Figure S5. Linear presentation of tracrRNA secondary structure.** The alignments of tracrRNAs from 10 selected VK_V1 sequences (upper) and all available VK_V2 sequences (lower) are shown below the arcs connecting predicted interacting nucleotides. The coloring of these arcs matches the respective stem-loops as indicated in the legend. Presumed crRNA binding regions are marked by red vertical bars and repeat regions by gray bars. TracrRNA regions are marked with blue lines. This figure extends **Fig. 3**.

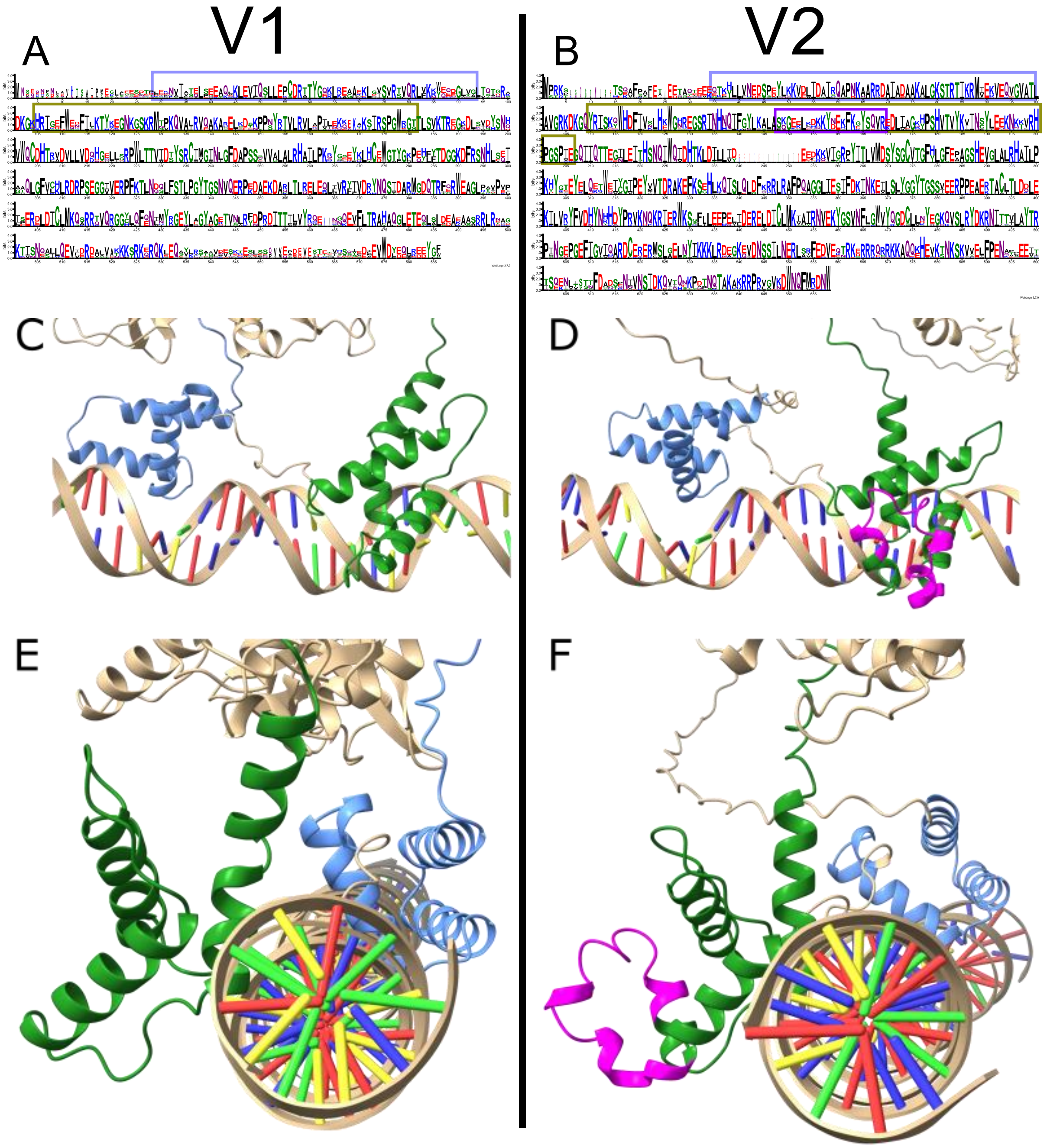


**Supplemental Figure S6. TnsB alignment and Alphafold N-terminal structure of V1 and V2 systems in comparison. (A)** Alignment of 9 related V1 TnsB sequences and **B.** 14 V2 TnsB sequences. The Alphafold N-terminal structures from TnsB of *Scytonema hoffmani* (WP_084763316.1) **(C, E)** and *Scytonema* sp. NIES-4073 (BAZ25524.1) **(D, F)** are shown with DNA containing IS elements from the related CAST systems. The same structures are depicted axial **(C, D)** and lateral **(E, F)** from the DNA strand. The DNA binding unit Iβ is colored in blue and Iγ is colored in green. The additional protein region of V2 TnsB is colored in violet. This figure extends **Fig. S2**. Additional information in **Dataset 7** and **8.**

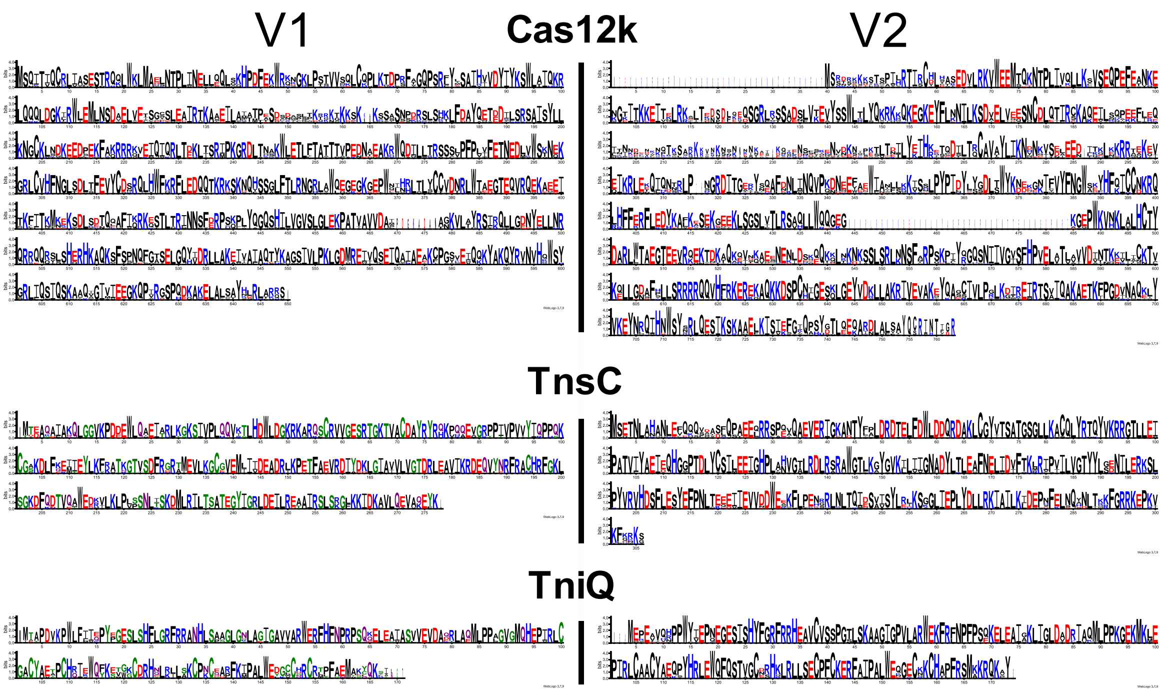


**Supplemental Figure S7. Alignment of Cas12k, TnsC and TniQ proteins of selected V1 CAST systems and all V2 systems.** The proteins for V1 systems from selected phylogenetic groups marked in their respective phylogenetic trees (**Fig. 2**, **S1**, **S3)** and only non-truncated sequences were used (10 proteins for Cas12k, 9 proteins for TnsC and TniQ). This figure extends **Fig. 2**, **Fig. S1** and **S3**.

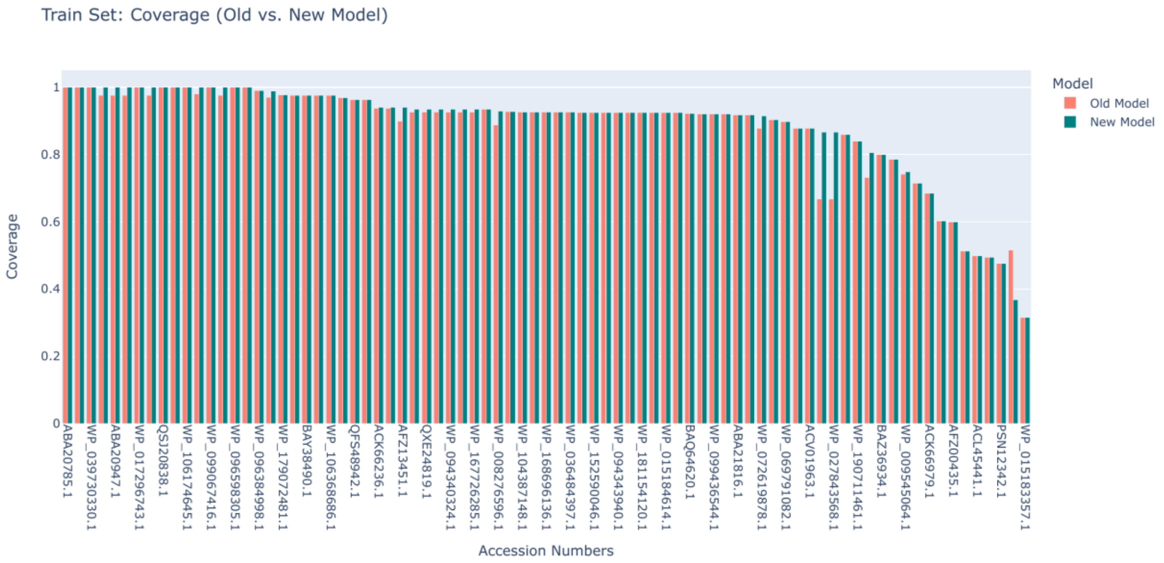


**Supplemental Figure S8. Training set coverage of CRISPRtracrRNA v.1.1.2.** Following the training of the new model, it was imperative to evaluate its performance to ensure that it did not deteriorate compared to the previous model. To achieve this, we assessed the model's coverage on the training dataset, quantifying its ability to retain relevant structure.

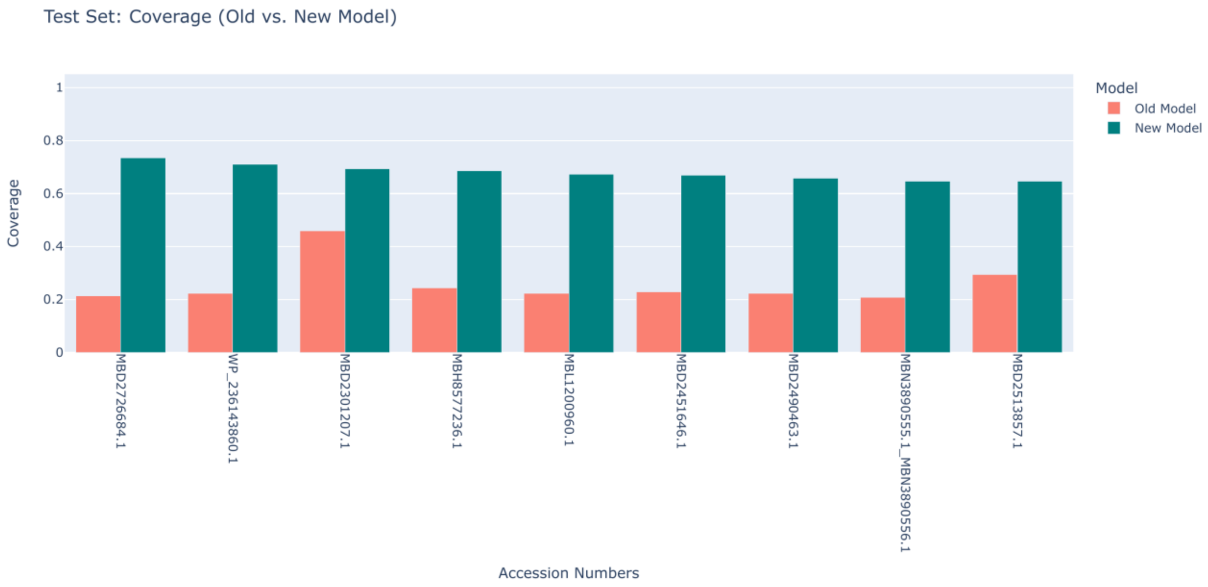


**Supplemental Figure S9. Test set coverage of CRISPRtracrRNA v.1.1.2.** The coverage level of the new model significantly improved on the test set compared to the old model. The new model consistently achieved approximately 75% coverage, whereas the old model exhibited an average coverage slightly above 20%. This substantial increase in coverage demonstrates the enhanced capability of the new model in capturing relevant structures.

### Supplementary Table

**Table S1. List of oligonucleotides used in this work.**

| Primer | Sequence | Purpose |
|---|---|---|
| Nblot_CRISPR_NIES 4073_fwd | TAATACGACTCACTATAGGGCCC AAAAGAAATGGCTTTCAACC | Northern blot probe (CRISPR array) via PCR with NIES-4073 genomic DNA as template |
| Nblot_CRISPR_NIES 4073_rev | TTCTCCACTAAACCGACGCC | |
| Nblot_tracrRNA_NIES 4073_fwd | TGCTCTTTTGAGCCTTAACTG | Northern blot probe (tracrRNA) via PCR with NIES-4073 genomic DNA as template |
| Nblot_tracrRNA_NIES 4073_rev | TAATACGACTCACTATAGGGACC CAAGCAATTACCGCTTAC | |
